# Cell-intrinsic IL4R alpha independence of large intestinal RELMα+ Ym1+ macrophages

**DOI:** 10.1101/2020.12.17.423033

**Authors:** Ruth Forman, Larisa Logunova, Hannah Smith, Kelly Wemyss, Iris Mair, Louis Boon, Judith E. Allen, Werner Muller, Joanne L. Pennock, Kathryn J. Else

**Author notes:** Correspondence to Ruth Forman and Kathryn J. Else.

## Abstract

The balance of pro-inflammatory and anti-inflammatory macrophages is critically important in enabling the development and resolution of inflammatory responses. Anti-inflammatory macrophages have been shown to be activated by IL4 and/or IL13 via the IL4Rα. In the context of type 2 immunity, anti-inflammatory macrophages have been defined by the expression of the signature markers RELMα, CD206 and Ym1, associated with activation of macrophages via the IL4Rα. Despite a breadth of inflammatory pathologies associated with the large intestine, many of which feature unbalanced macrophage activation states, little is known about how large intestinal macrophages are activated. Here, we address this important knowledge gap by using a *Trichuris muris* infection model of resolving type 2 intestinal inflammation, in combination with transgenic mice (IL4Rαfl/fl.CX3CR1Cre) and IL4Rα-deficient/wild-type mixed bone marrow chimaeras. These models allowed us to interrogate the role of IL4/IL13 in macrophage activation driven by inflammation of the large intestine. We make the unexpected finding that education of large intestinal macrophages towards a RELMα and Ym1 expressing cell type during type 2 inflammation, does not require IL4Rα expression on macrophages. Thus, upregulation of RELMα and Ym1 post infection is independent of macrophage IL4Rα expression. Further, this independence is maintained even when the mice are treated with anti-IFNγ antibody to create a strongly polarised Th2 environment. In contrast to RELMα and Ym1, PD-L2 expression on macrophages post infection was dependent on IL4Rα signalling in the macrophages. These data challenge existing paradigms, evidencing that expression of RELMα and Ym1 by macrophages, typically regarded as having anti-inflammatory functions, do not always rely on IL4/IL13.

## INTRODUCTION

The activation of macrophages is a fundamental process in tissue homeostasis, as well as in chronic and resolving inflammation. However, a full understanding of the *in vivo* activating signals, which enable a balance of macrophage subpopulations to emerge and regulate inflammatory processes, is lacking. This is essential if we are to understand the mechanisms of chronic inflammation and develop targeted therapies to improve disease outcome (1).

Distinct macrophage polarization was first described by Gordon and colleagues in the context of classical activation, driven by IFNγ and LPS; and alternative activation, driven by IL4 and IL13 (2, 3). It later became clear that many different stimuli could lead to an anti-inflammatory macrophage phenotype, collectively referred to as M2. The term M2 thus embraces a wide spectrum of antiinflammatory, alternatively activated macrophage states and consequently has lost some utility. Murray and colleagues (4) attempted to bring clarity to M2 nomenclature describing a set of markers associated with alternatively activated macrophages activated *in vitro* by IL4. These macrophages, coined M(IL4), are identified as RELMα^+^ Ym1^+^CD206^+^ and PD-L2^+^ macrophages. Many laboratories have demonstrated that these markers are also IL4Rα dependent *in vivo* and these have become reliable indicators of M(IL4), especially in the context of type 2 inflammation.

More recently, during steady state, IL4Rα-independent pathways of Ym1 and RELMα expression have been described in lung macrophages (5). Such steady state IL4Rα–independence was subsequently shown not to be limited to the lung, with both IL4Rα and the downstream transcription factor STAT6 dispensable for RELMα expression in steady state peritoneal macrophages (6). Beyond the steady state, there is also a precedent in the literature for the induction of RELMα on macrophages to be partially IL4Rα independent in the context of lung and liver inflammation, following *Nippostrongylus brasiliensis* and *Schistosoma mansoni* infection respectively (5, 7). Moreover, using the global IL-4Rα deficient mouse, an IL4Rα-independent macrophage activation resulting in equivalent expression of RELMα, Ym1 and Arg1 has been observed in the peritoneal cavity driven by lactate (8). Similarly, tumour associated macrophages in the global IL-4Rα deficient mouse have been shown to express equivalent RNA levels of RELMα and Arg1 driven by hypoxia (9). Looking beyond tumour, liver, lung and peritoneal macrophages, it remains unknown whether intestinal macrophages depend on IL4Rα signalling for their education. Attempts to address the importance of the IL4Rα in the context of large intestinal macrophages have been limited by the Cre driver employed to delete the IL4Rα from macrophages and the markers used to define the M2 phenotype. Thus, recent work using oxazalone-induced colitis in IL4Rαfl.LysM cre mice demonstrated no change in total Arg-1 mRNA expression in the intestine post infection (10). However, no macrophage specific expression of M2 markers was reported and the interpretation of this data is confounded by the known inefficiencies in LysM cre mediated deletion of the IL4Rα on macrophages during inflammation (11). Thus, there remains no clear exploration of the role of IL4Rα in the education of large intestinal macrophages during either steady state or in an inflamed gut environment. This represents a surprising gap in our knowledge, given that it is essential information underpinning studies striving to enrich for anti-inflammatory macrophages in pro-inflammatory settings such as inflammatory bowel disease.

The ability to control inflammation in the large intestine is vitally important, as a breach in the epithelial lining can expose the host immune system to potentially lethal gut microbes. Macrophages are the most common mononuclear phagocyte in the healthy intestinal lamina propria and act as vital immune sentinels, able to sense and respond to changes within the gut. Macrophage dysfunction results in an excessive inflammatory response and a failure to resolve inflammation (12). In Inflammatory Bowel Disease (IBD) and animal models of IBD, an excess of pro-inflammatory macrophages and altered monocyte-macrophage differentiation are observed promoting the release of pro-inflammatory cytokines (13–15). In contrast, anti-inflammatory macrophages have been proposed to alleviate severe IBD development (16–18). Thus enabling intestinal macrophage education towards an anti-inflammatory phenotype has been considered as a target for potential therapeutic intervention (19, 20) if the educating signals could be defined.

In contrast to IBD, where pro-inflammatory macrophages predominate, anti-inflammatory macrophages are a feature of many helminth infections (21). We have recently defined an *in vivo* model of resolving intestinal inflammation, involving infection of mice with the gut-dwelling nematode parasite *Trichuris muris (T. muris).* Expulsion of the parasite requires the generation of a Th2 immune response, with IL4 and IL13 essential in mediating resistance. RELMα^+^ macrophages emerge in the gut in a predictable manner (22) and, post-expulsion, intestinal inflammation resolves and the gut architecture is restored. Macrophages dominate the lamina propria during the inflammatory response and its subsequent resolution following worm expulsion (22, 23). The RELMα^+^macrophage is most abundant post-infection, during the resolution of inflammation. Thus, this robust *in vivo* model enables definition of the signals driving the emergence of RELMα^+^macrophages during and after intestinal inflammation.

We set out to evaluate the role of IL4Rα in educating macrophages in the inflamed type 2 polarised large intestine, given the recognised importance of IL-4 and IL-13 in promoting anti-inflammatory macrophages in other inflamed tissues (4, 5, 8, 24, 25). To our surprise, using a combination of *in vivo* approaches, including transgenic mice (IL4Rαfl/fl.CX3CR1Cre) and mixed bone marrow chimaeras, we found that during inflammation, IL4Rα on large intestinal macrophages was completely dispensable for the emergence of RELMα^+^Ym1^+^ macrophages. This work is vital to inform strategies treating excessive, chronic intestinal inflammation as it highlights treatments which target the IL4R to promote anti-inflammatory macrophages are unlikely to be efficacious. Our work also demonstrates the need to understand more about macrophage biology *in vivo* in physiological contexts, and to corroborate *in vitro* observations with *in vivo* evidence investigating macrophages at sites of inflammation.

## RESULTS AND DISCUSSION

### Intestinal inflammation associated with a *Trichuris muris* infection drives development of RELMα^+^macrophages *in vivo*

Infection with the gut dwelling nematode parasite *T. muris* is associated with the development of antiinflammatory RELMα^+^ macrophages (22). To characterise these macrophages we performed bulk RNA seq on monocytes and macrophages isolated from the caecum and colon of infected C57BL/6 mice. Monocyte and macrophage populations were defined based on the previously described monocyte/macrophage waterfall where Ly6C^hi^ blood monocytes enter the intestine (P1), progressively upregulate surface MHC II expression (P2) and down regulate Ly6C expression to become mature gut macrophages (P3) (26–29). Cells were sorted using FACS and isolated into two subsets based on surface expression of the marker CD206, chosen based on good co-expression with intracellular RELMα post *Trichuris* infection (data not shown). Principal component analysis (PCA) analyses of the two monocyte / macrophage subsets revealed 58% of the sample variation to be attributed to expression of CD206. Importantly, analysis of the top 20 genes up- and down-regulated between the 2 macrophage populations revealed that the CD206^+^ macrophage profile associated with genes such as *Retnla, Mrc1, Ccl8, CD163, Ccl6,* and *Gas6* (30) (Supp Fig 1a,b). Because the transcriptional landscape fits the canonical anti-inflammatory macrophage phenotype (24, 30), this data strongly supported using *T. muris* infection as a model for understanding macrophage activation towards an anti-inflammatory phenotype in the large intestine.

### IL4Rαfl/fl.CX3CR1Cre+ macrophages are un-responsive to IL4 stimulation

To determine the *in vivo* role of IL4/IL13 in macrophage education we crossed CX3CR1Cre mice (31) with IL4Rαfl/fl mice (32) to generate IL4Rαfl/fl.CX3CR1Cre mice. Macrophages from these IL4Rαfl/fl.CX3CR1Cre+ mice, (hereafter referred to as Cre+), lack the IL4Rα chain, and therefore are unable to respond to IL4 and IL13. In contrast, littermate controls (IL4Rαfl/fl. CX3CR1Cre-) remain IL4/IL13 responsive. The Cre driver was chosen as it is known to efficiently delete floxed alleles in all intestinal macrophages (31). This is important, as historically, determining the function of genes in intestinal macrophages has been confounded by inefficacies in deletion of floxed genes in the gut when using the LysM Cre driver (11). The inefficient deletion of genes by LysM Cre is observed in subpopulations of macrophages which express low levels of Lyz2. Therefore, whilst LysM Cre is able to efficiently delete IL4Rα under steady state conditions, during inflammation a large proportion of macrophages retain IL4Rα expression (11, 33). In order to confirm the efficient deletion of the IL4Rα in CX3CR1Cre^+^ large intestinal macrophages, we injected mice intraperitoneally with a recombinant IL4 complex (IL4c). Under uninflamed steady state conditions, RELMα expression in naïve macrophages from the peritoneal cavity (Fig 1a,b), gut (Fig 1c,d) and liver (Fig 1e,f) all showed no dependency on IL4Rα in keeping with the literature from lung (5) and peritoneal macrophages (6). Using the IL4Rαfl/fl.CX3CR1Cre mice, and in accordance with data previously reported with global IL4Rα deficient mice (34), IL4c drove an IL4Rα-dependent activation of F4/80^hi^ macrophages within the peritoneal cavity, with an upregulation of RELMα only detectable in the Cre-mice (Fig 1a,b). Importantly, this is phenocopied in the large intestinal macrophages where IL4c drove strong RELMα expression in the Cre-but not Cre+ mice (Fig 1c,d). Likewise, in the liver, Kupffer-cell driven expression of RELMα was dependent on the absence of Cre (Fig 1e,f). Our results thus far therefore support the robust deletion of the IL4Rα from large intestinal macrophages in the IL4Rαfl/fl.CX3CR1Cre mouse, with IL4Rα signalling required for the induction of RELMα^+^ macrophages. To corroborate these data further, IL4Rα expression on large intestinal macrophages was determined at d35 post infection with *T. muris,* to control for the possibility of a selective outgrowth of a small population of Cre+ macrophages with incomplete deletion of the IL4Rα. These data revealed low expression of the IL4Rα on colonic macrophages in Cre-mice which was absent in the Cre+ mice (Supp Fig 2a). Generation of bone marrow macrophages from IL4Rαfl/fl.CX3CR1Cre mice allowed a more comprehensive profiling of these macrophages and confirmed that macrophages generated from Cre+ mice were unable to express the IL4Rα even following stimulation with the strong IL4Rα driver IL6 (35) (Supp Fig 2b). Additionally, bone marrow macrophages from Cre+ mice were unable to upregulate the canonical anti-inflammatory macrophage markers CD206, RELMα and Ym1 following exposure to IL4 (Supp Fig 2c-e).

**Figure 1:**
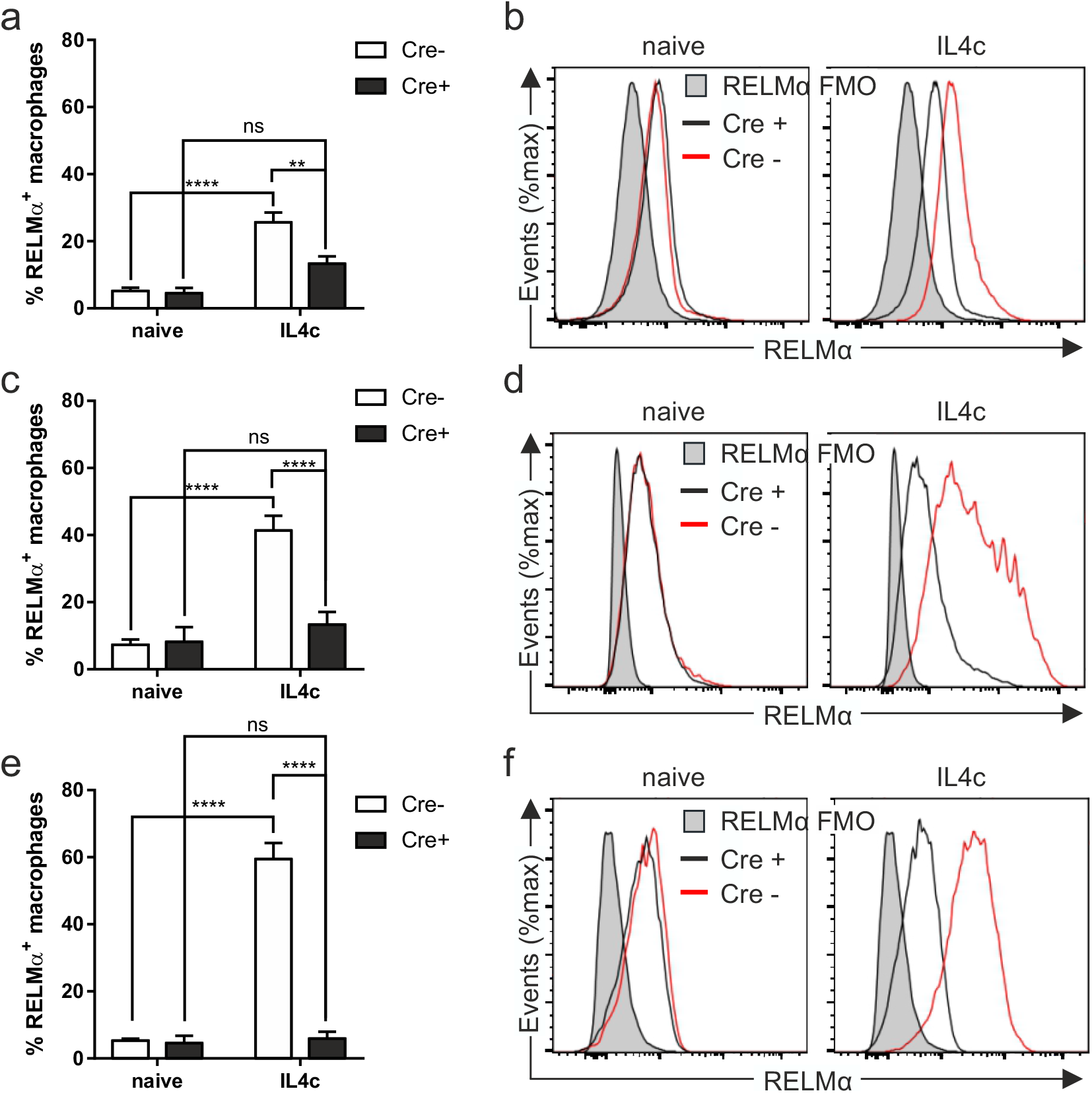
IL4R^fl/fl^ CX3CR1Cre^+^ macrophages are unresponsive to IL4 signalling. IL4R^fl/fl^ CX3CR1Cre^+^(Cre^+^) or IL4R^fl/fl^ CX3CR1Cre^-^ (Cre^-^) mice were injected intraperitoneally with recombinant IL4 complex and expression of RELMα analysed in the (a) peritoneal cavity, (c) colon and (e) liver 48hours later. Representative histograms show RELMα expression in the Cre^-^ (red) or Cre^+^ (black) mice in the (b) peritoneal cavity, (d) colon and (f) liver compared to the FMO control. Data are combined from two independent experiments (naïve; n=3-7, d18; n=6). Error bars show means ± SEM. Statistical comparisons were performed with a two-way ANOVA with post-hoc Bonferroni’s multiple comparison test where appropriate: \**p*<0.05, \*\**p*<0.01, \*\*\**p*<0.001, \*\*\*\**p* < 0.0001 indicates significance as indicated.

### IL4Rαfl/fl.CX3CR1Cre mice respond to *T. muris* infection with normal kinetics

Given that the IL4Rαfl/fl.CX3CR1Cre mouse is a novel transgenic mouse we characterised the expulsion kinetics and immune response to *T. muris* post-infection. We saw no differences between the worm burdens of Cre+ and Cre-mice (Fig 2a) or the quality of the immune response. Thus, as is typical of mice on a C57BL/6 background (22), infection drove a mixed IgG1/IgG2c response (Fig 2b-g). Furthermore, a mixed cytokine response was seen in the draining mesenteric lymph nodes (MLN), with MLN cells making elevated levels of IFNγ, IL4, IL13 and TNF compared to naïve mice (Fig 2h-k), but with no significant differences between genotype. As expected, post infection in both genotypes we saw a significant crypt hyperplasia (Fig 2l,m) in the descending proximal colon of the mice, which remained elevated after worm expulsion (day 35 post infection). This was accompanied by a significant goblet cell hyperplasia at d18 post infection (Fig 2l,n) resolving to normal levels by day 35 (Fig 2l,n).

**Figure 2:**
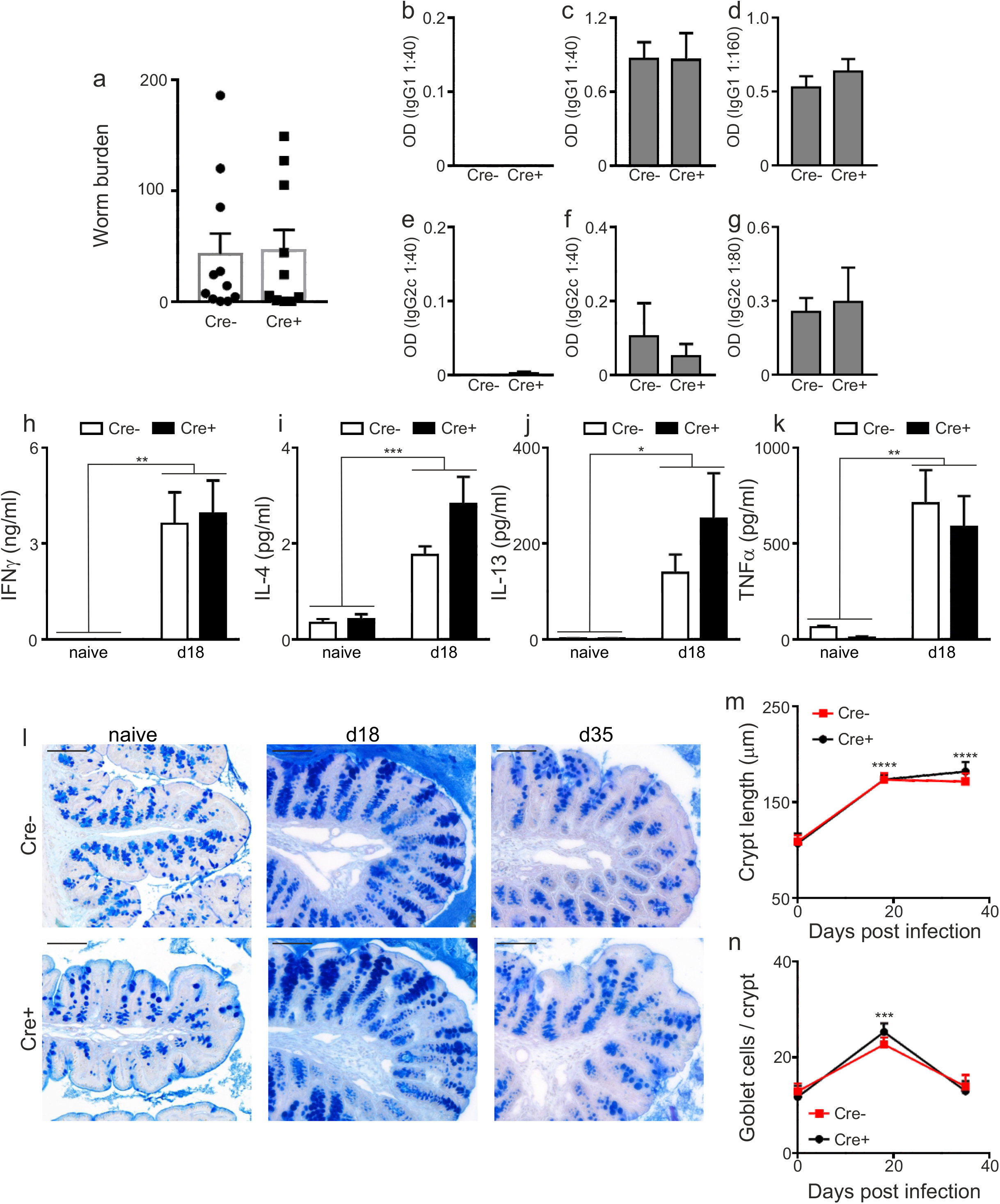
The kinetics of *T. muris* infection is unaltered in the absence of IL4Rα on macrophages. IL4R^fl/fl^ CX3CR1Cre^+^ (Cre+) or IL4R^fl/fl^ CX3CR1Cre^-^ (Cre^-^) mice were infected with 200 *T. muris* eggs and (a) worm burden assessed in the colon and caecum at d18 post infection (n=10-11, pooled from 2 independent experiments). Parasite-specific IgG1 from (b) naïve (n=6-7, pooled from 2 independent experiments), (c) d18 post infection (n=6, representative of 3 independent experiments) and (d) d35 post infection (n=5-7, pooled from 2 independent experiments) and IgG2c production from (e) naïve, (f) d18 post infection and (g) d35 post infection. Mesenteric lymph node (MLN) cell (h) IFNg, (i), IL4, (j) IL13 and (k) TNF profiles in naïve mice (n=3-4, representative of 2 independent experiments) and at d18 post infection (n=6, representative of 3 independent experiments) cultured with *T. muris* E/S (50μg/ml) for 48h. Supernatants analysed using cytometric bead array. (l) Representative photographs of periodic acid/Schiff’s staining in colonic tissue with quantification of (m) crypt lengths and (n) goblet cell numbers in colonic tissue of naïve (n=4-6, 2 independent experiments), d18 (n=11-12, 2 independent experiments) and d35 (n=5, 2 independent experiments) post infection mice. Scale bar indicates 100μm. Error bars show means ± SEM. Statistical comparisons were performed with a two-way ANOVA with post-hoc Bonferroni’s multiple comparison test where appropriate: \**p*<0.05, \*\**p*<0.01, \*\*\**p*<0.001, \*\*\*\**p*< 0.0001 indicates significance compared to naïve control.

Together these data demonstrate that our subsequent analyses of *in vivo* macrophage-drivers in the IL4Rαfl/fl.CX3CR1Cre mouse are not confounded by differences in expulsion kinetics or immune response between genotypes.

### Intestinal macrophages are educated towards RELMα^+^macrophages in the absence of the IL4Rα subunit

To determine the role of the IL4Rα in shaping the colonic macrophage pool under inflammatory settings, we characterised the large intestinal macrophage population in naïve mice and at two key time points post infection – d18 during active inflammation and at d35 during the resolution of inflammation. Analyses of macrophages within the P1-P3 gates (26–29), as well as the recently defined Tim4^-^CD4^-^, Tim4^-^CD4^+^ and Tim4^+^CD4^+^ gates within the mature (P3) macrophage population (36), revealed equivalent population shifts in both Cre+ and Cre-mice over the time course of infection (Fig 3a-c). Thus, as anticipated, during active inflammation, we observed a significant increase in the number of recruited monocytes into the intestine (P1, P2), accompanied by an enrichment in the Tim4^-^CD4^-^ population within P3 gate, in keeping with the monocyte-derived origin of this population (36). Based on our IL4c experiments, we hypothesized that RELMα^+^ macrophages in the Cre+ mice would be altered. However, analysis of RELMα expression within the monocyte/macrophage waterfall during both active (d18) and resolving (d35) inflammation revealed no deficit in RELMα expression in the IL4Rα null (Cre+) macrophages (Fig 3d,e), with both Cre+ and Cre-mice showing significant increases in RELMα expression post infection. Moreover, macrophage RELMα expression was selectively enriched in the more mature (Tim4+) P3 macrophage subsets. Thus, enhanced RELMα expression was observed in the Tim4^+^CD4^-^ and Tim4^+^CD4^+^ populations compared to the Tim4^-^CD4^-^ population. However the expression of RELMα by Tim4+ macrophages was seen in both the Cre+ and Cre-mice and thus independent of the IL4Rα (Fig 3f). Immunohistochemical analyses of RELMα and CD68 coexpression within the colon revealed analogous results, with no deficiency in RELMα expression in the Cre+ mice and no alterations in the spatial location of the RELMα^+^CD68^+^ macrophages (Fig 3g). We also analysed the expression of other signature markers associated with anti-inflammatory macrophages; specifically PD-L2, CD206 and Ym1, in addition to RELMα. Interestingly, although the expression of CD206 did not change post infection (Fig 3h), Ym1 (Fig 3i) was also induced *in vivo* post infection independently of IL4Rα. In contrast, induction of PD-L2 expression was IL4Rα-dependent. Thus PD-L2 expression was induced post infection in the Cre-mice but the % PD-L2^+^ macrophages evident in the Cre+ mice post infection was significantly reduced (Fig 3j). Therefore, our data reveal that the emergence of the large intestinal RELMα^+^Ym1^+^ macrophage is independent of IL4/IL13.

**Figure 3:**
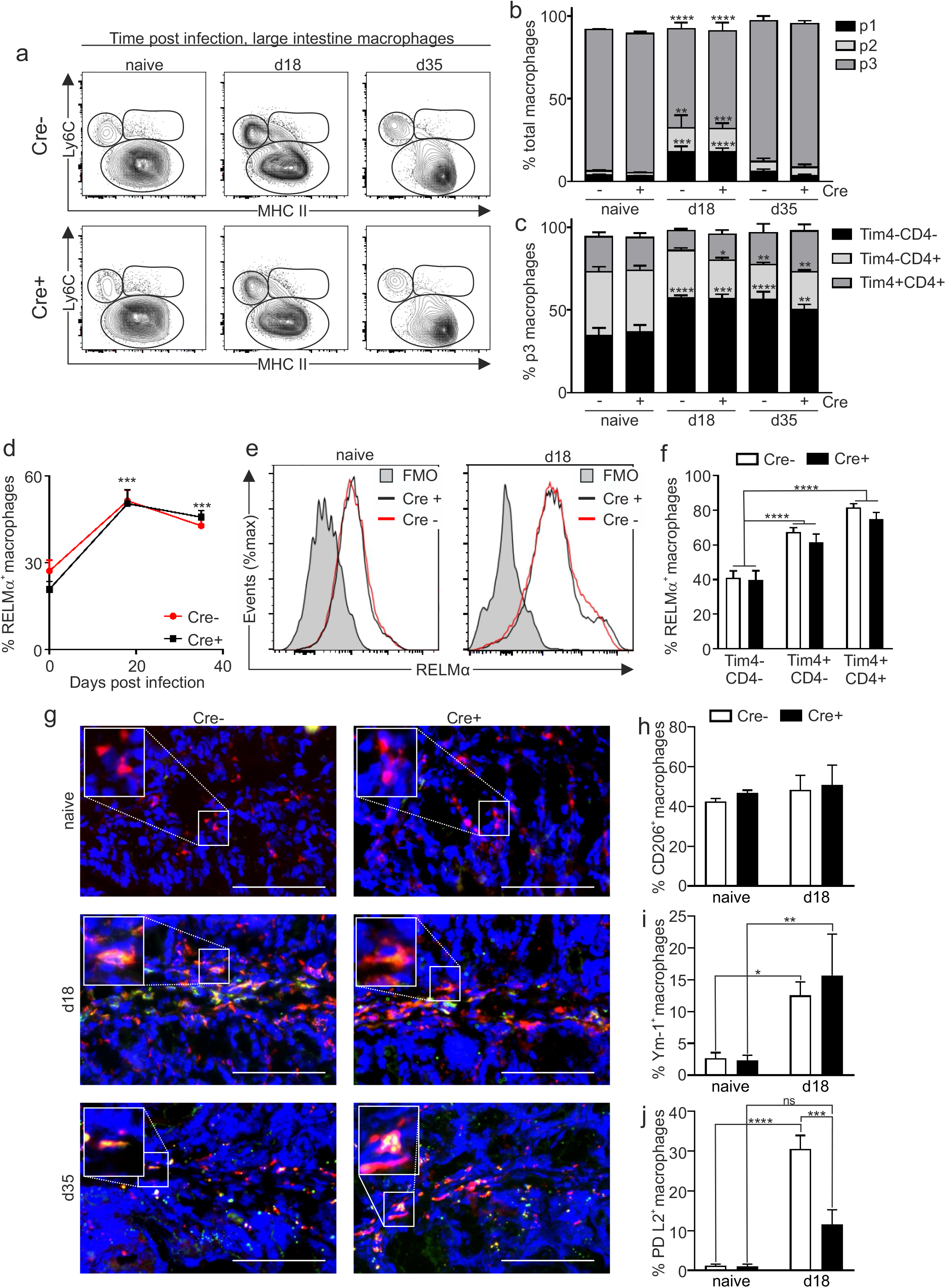
Intestinal RELMα^+^macrophages develop independently of IL4Rα expression following *T. muris* infection. Gut macrophages (gated as CD45^+^CD11b^+^CD11c^low/int^SiglecF^-^Ly6G^-^) were divided into P1 monocyte (CD64^-^Ly6C^hi^MHCII^-^), P2 transitioning monocyte (Ly6C^+^MHCII^+^) and P3 macrophage (CD64^+^Ly6C^-^MHCII^+^) compartments in the colon and caecum. (a) Representative plots illustrating the P1-3 gates in Cre^-^ and Cre^+^ mice at the defined time points. (b) Proportion of macrophages within the P1-P3 and (c) Tim4^-^CD4^-^, Tim4^-^CD4^+^ and Tim4^+^CD4^+^ gates in naïve mice and post infection with *T. muris;* naïve (n=6-7, pooled from 2 independent experiments), d18 post infection (n=3, representative of 3 independent experiments) and d35 post infection (n=5-7, pooled from 2 independent experiments). Expression of the marker RELMα (d) within the total monocyte/macrophage (P1-3) population and (e) representative histograms showing RELMα expression in the Cre- (red) and Cre+ (black) naïve mice and at d18 post infection with the relevant FMO control (shaded grey) naïve (n=6-7, pooled from 2 independent experiments), d18 post infection (n=7-9, pooled from 2 independent experiments) and d35 post infection (n=5-7, pooled from 2 independent experiments. (f) Expression of RELMα in P3 within the Tim4^-^CD4^-^, Tim4^-^CD4^+^and Tim4^+^CD4^+^ gates at d18 post infection (n=7-9, pooled from 2 independent experiments). (g) Representative staining of colonic tissues stained for CD68 (Red), RELMα (Green) and counterstained with DAPI (Blue), Scale bar shows 100μm. Analysis of additional markers (h) CD206, (i) Ym1 and (j) PD-L2 in naïve mice and at d18 post infection (n=4-7; representative of 2 independent experiments) Error bars show means ± SEM. Statistical comparisons were performed with a two-way ANOVA with post-hoc Bonferroni’s multiple comparison test where applicable or by student’s *t*-test. Significance is shown compared to naïve control \**p*<0.05, \*\**p*<0.01, \*\*\**p*<0.001, \*\*\*\**p*< 0.0001 or as indicated.

Further, our data reveal a discord in the regulation of the anti-inflammatory macrophage marker PD-L2 in comparison to Ym1 and RELMα with a small IL4-dependent population of large intestinal macrophages present, characterised by the expression of PD-L2.

### IL4Rα deficient macrophages in a mixed bone marrow chimera express CD206, RELMα and Ym1 at levels equivalent to IL4Rα sufficient macrophages

Given our unexpected observations, we corroborated our findings using an independent experimental approach taking advantage of the IL4Rα global knockout mouse. However, as IL4Rα global deficient mice are unable to expel *Trichuris* as they do not develop a Th2 immune response, we employed a mixed bone marrow chimera approach. Thus, CD45.1/CD45.2 recipient mice were lethally irradiated and reconstituted with a 50:50 bone marrow mix from CD45.1 (IL4Rα+/+; WT) mice and CD45.2 IL4Rα global deficient mice (IL4Rα-/-) enabling IL4Rα-dependency to be determined through analysis of CD45.1 expression (Fig 4a). At 8 weeks after irradiation mice were tail bled to analyse baseline chimerism, and mice were subsequently infected with *T. muris.* Analysis of blood chimerism revealed that blood-borne monocytes and other populations (CD4^+^ T cells, granulocytes) exhibited roughly equal proportions of CD45.1 and CD45.2 cells after reconstitution and these proportions were not altered by *T. muris* infection (Supp Fig 3a). This is in keeping with previously published data showing IL4Rα signalling is not required for steady state survival, or survival of blood monocytes following helminth infection (34, 37). Additional analysis post-infection in both the spleen (Supp Fig 3b) and MLNs (Supp Fig 3e) revealed no preference for CD45.1 or CD45.2 cells. Re-stimulation of MLNs with PMA and Ionomycin following infection demonstrated an equal contribution of both CD45.1 and CD45.2 cells to CD4^+^ and CD8^+^ T cell derived IFNγ and IL13 production (Supp Fig 3c-e) suggesting that, despite lacking the IL4Rα, CD45.2 cells are able to produce both Th1 and Th2 cytokines. This may reflect the ability of these cells to respond to other Th2-driving stimuli, for example IL33 or CCL-2 (38) and is consistent with data showing IL4 signalling is not required for the initial generation of Th2 cells in the lymph node (39). The chimeric mice were able to respond in a typical manner to a *T. muris* infection mounting a mixed IgG1/IgG2c response (Supp Fig 3f,g) and a mixed cytokine response in the draining MLNs (Sup Fig 2h-m). Interestingly, and in contrast to our blood monocyte data, we saw a selection for IL4Rα-/- monocytes in the colon, with a slight but significant decrease in cells derived from the CD45.1 (WT) compartment compared to in the bloodstream (Fig 4b,c). This occurred independently of infection. Importantly, infection with *Trichuris* drove an influx of monocytes into the colon as previously observed in the IL4Rαfl/fl.CX3CR1Cre mice with an increase in the P1 subset at day 18 post infection (Fig 4d). Analysis of the chimerism between the different monocyte and macrophage subsets in the colon again revealed a decrease in the monocytes derived from the wild type donor (CD45.1) in the intermediary P2 transiting monocyte subset in both the naïve and infected mice, however this did not appear to have a subsequent effect on the chimerism of the resident P3 macrophages (Fig 4e,f). Importantly, analysis of the RELMα^+^ population showed no effect of donor, demonstrating the induction of macrophage RELMα post infection with *T. muris* is independent of IL4Rα expression (Fig 4g,i). Analogous results were also observed for Ym1 and CD206 expression (data not shown). As in the IL4Rαfl/flCX3CR1Cre mice experiments, in our mixed bone marrow chimeras, PD-L2 expression was again dependent on the presence of IL4Rα (Fig 4h,i). As PD-L2 expression has been described to be exclusively observed on macrophages derived from infiltrating monocytes and not tissue resident tissue macrophages within the peritoneal cavity (40) we analysed the PD-L2+ macrophages to determine where they fell within the P1, P2 and P3 gates post infection, to determine if the altered chimerism in the P2 monocyte subset was influencing the overall PD-L2 expression. Our data shows that, in contrast to macrophages within the peritoneal cavity, PD-L2+ macrophages are predominantly found in the more mature ‘P3’ macrophage subset within the intestine (Fig 4j). The IL4Rα^+^ independent induction of RELMα and Ym1 on large intestinal macrophages is in direct contrast to seen with cavity macrophages. Thus, mixed bone marrow chimeras utilising IL4Rα deficient and sufficient bone marrow have shown that pleural cavity and peritoneal cavity macrophages express RELMα and Ym1 in an IL4Rα dependent way following infection with *Litomosoides sigmodontis* and *Heligmosomoides polygrus* respectively; the authors did not report analysis of PD-L2 expression in these studies (34). Further, in the *Litomosoides* infection model at d16 (but not d10) post infection IL4Rα expression bestowed a selective advantage specifically on the F4/80^hi^ pleural macrophage population which was proposed to reflect enhanced entry of the IL4Rα^+^ cells into the cell cycle. Thus, the differences in IL4/IL13 dependency of macrophages seen between these studies and our data may be due in part to proliferative capacity of the macrophages analysed, with the macrophage population following *T. muris* infection not undergoing the vast proliferation seen following an infection with *Litosomoides sigmodontis* (22, 41). Thus the selective advantage provided by the presence of the IL4Rα is not observed.

**Figure 4:**
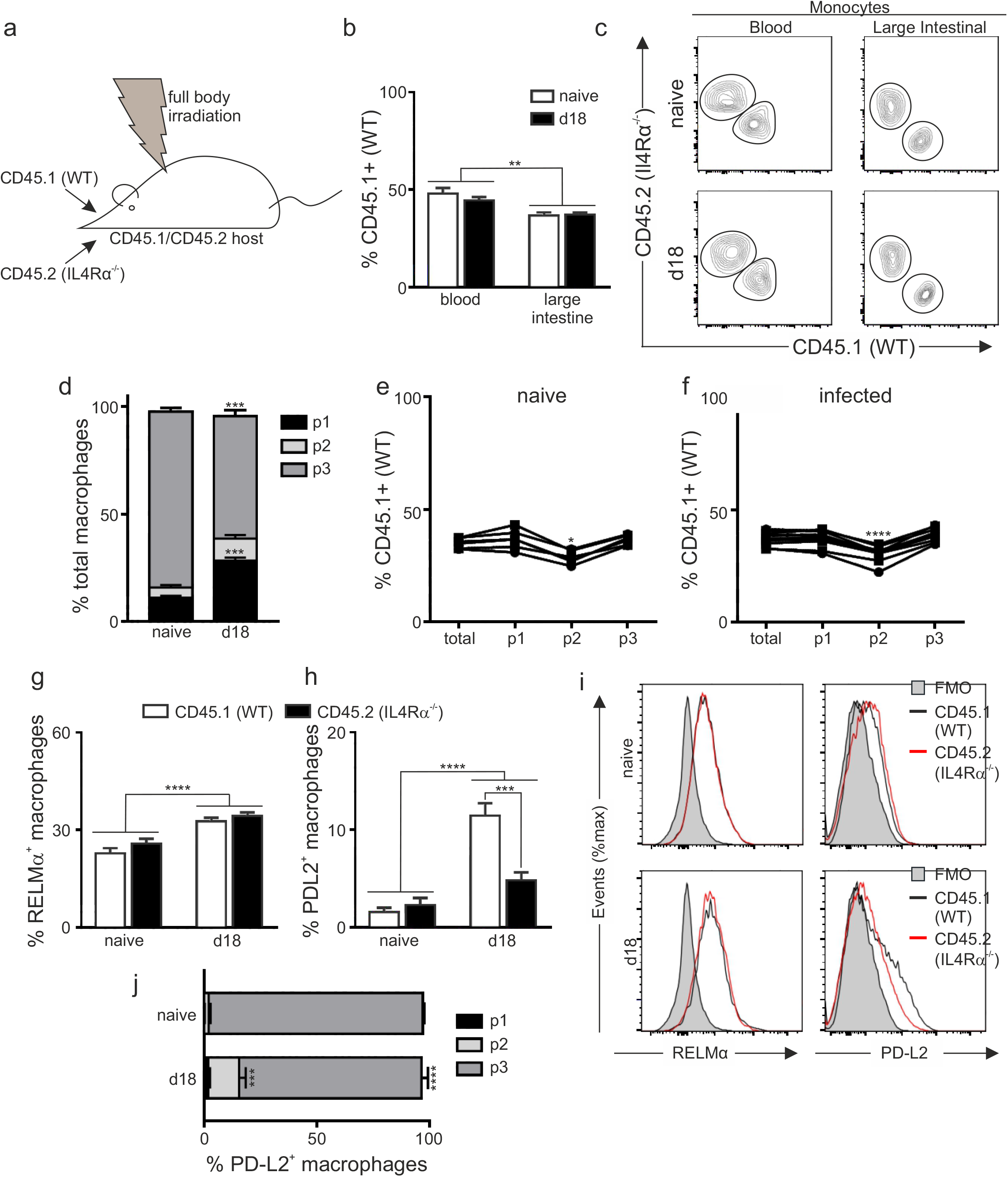
IL4Rα signalling is dispensable for RELMα^+^macrophage development in the gut following *T. muris* infection of mixed bone marrow chimeras. (a) Mixed bone marrow chimeras were generated by lethally irradiating C57BL/6 IL4Rα^+/+^CD45.1^+^CD45.2^+^ mice and reconstitution with a 50:50 mix of IL4Rα^+/+^ CD45.1 (WT) and IL4Rα^-/-^ CD45.1nullCD45.2^+/+^ congenic bone marrow cells; mice were left for 8 weeks to reconstitute. Mice were left uninfected or infected with 200 *T. muris* eggs and (b) Quantification of chimerism in the blood and gut monocyte population in naïve mice and at d18 post infection. (c) Representative staining for CD45.1 and CD45.2 shows chimerism of the blood monocytes (CD115^+^) and gut monocyte (infiltrating P1 population) in naïve and infected mice. (d) Frequency of monocytes / macrophages with the P1-P3 waterfall in naïve and d18 post infection mice and the chimerism within these populations in (e) naïve and (f) infected mice. Frequency of the (g) RELMα and (h) PD-L2 expressing macrophages in the total P1-3 macrophage population from naïve and d18 post infection mice in the CD45.1 and CD45.2 populations. (i) Representative flow histograms are shown for naïve and d18 post infection mice. (j) Location within the P1-P3 waterfall of PD-L2^+^ monocytes / macrophages in naïve mice and at d18 post infection. Data are combined from two independent experiments (naïve n=6, d18 n=11). Error bars show means ± SEM. Statistical comparisons were performed as appropriate by repeated measure or standard two-way ANOVA with post-hoc Bonferroni’s multiple comparison test where applicable. Significance is shown compared to naïve control * *p*<0.05, \*\**p*<0.01, \*\*\**p*<0.001, \*\*\*\**p*< 0.0001 or as indicated.

Differences in IL4/IL13 dependency between gut and cavity macrophages may also reflect the length of tissue residency, with the F4/80^hi^ macrophages in serous cavities which respond to helminth infections being the resident macrophage population (41), and not derived from infiltrating monocytes as observed post-infection with *T. muris.* It is possible therefore that the monocyte-derived intestinal macrophage has a less stringent dependency on IL4/IL13 signalling in order to express RELMα than resident macrophages. Indeed, using Tim4+CD4+ expression as markers of length of residency in the gut, our data showed an increased expression of RELMα within the Tim4^-^CD4^+^ and Tim4^+^CD4^+^ intestinal P3 macrophage subsets compared to the Tim4^-^CD4^-^ population, however this too was independent of IL4Rα expression.

### Intestinal macrophages from IL4Rαfl/fl.CX3CR1Cre mice become RELMα^+^ during *T.muris* infection even following αIFNγ treatment

The recent reports of a partial IL4Rα independence of anti-inflammatory macrophages in tissues other than the large intestine is particularly evident prior to the development of highly polarised type 2 adaptive immune responses (5). Thus we sought to determine if the IL4Rα-independence of RELMα^+^macrophages education in the inflamed large intestine was due to the mixed Th1/Th2 environment present, as opposed to a highly polarised Th2 setting. In order to drive a highly polarised Th2 immune response, we treated IL4Rfl/fl.CX3CR1Cre mice with an αIFNγ antibody throughout the course of infection (Fig 5a). αIFNγ treatment during *T. muris* infection is a well-established methodology known to accelerate worm expulsion due to a shifting of the immune response towards a Th2 response (42, 43). There was no change in the distribution of macrophages within the P1-P3 macrophage waterfall in the αIFNγ treated mice (Fig 5b,c). However, both parasite-specific IgG1 and IgG2c levels in the serum were significantly reduced in αIFNγ treated mice, independently of IL4Rα expression, as typically seen in mice which expel the parasite very rapidly (Fig 5d,e) (44). Consistent with an enhanced Th2 response we saw a trend towards elevated serum IgE in αIFNγ depleted mice (Fig 5f). Importantly, although treatment with αIFNγ drove an increase in RELMα^+^ macrophages in the colon (Fig 5g) this occurred independently of IL4Rα expression on the macrophages. Interestingly, the PD-L2 dependence on IL4Rα expression appeared to be lost in the more strongly polarised environment suggesting that the contribution of IL4Rα signalling to PD-L2 expression may be less stringent in strong Th2 settings, although this may also reflect a return to steady state conditions due to the accelerated worm expulsion (Fig 5h).

**Figure 5:**
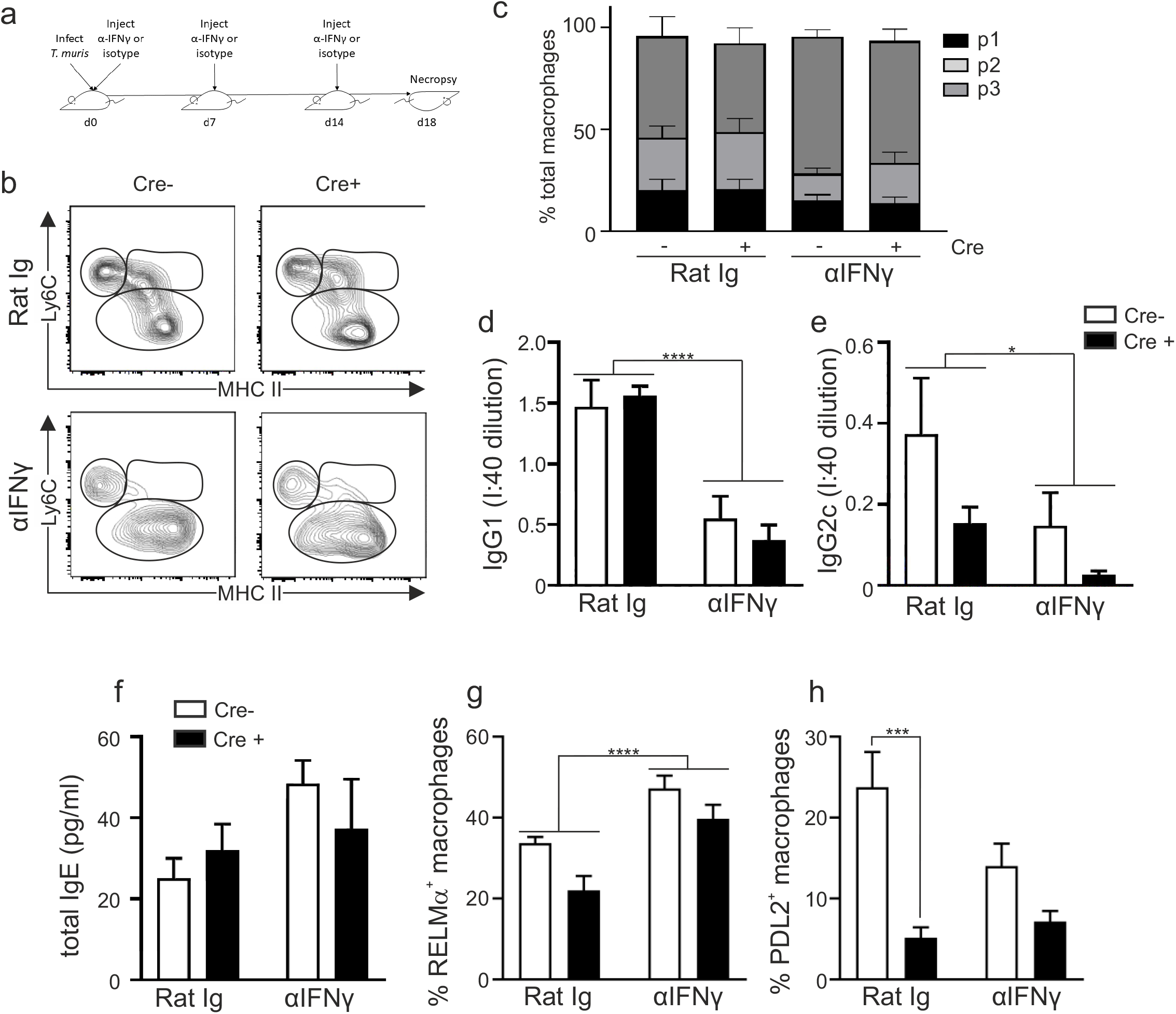
Intestinal RELMα^+^ macrophages develop independently of IL4Rα expression following αIFNγ treatment during *T. muris* infection. (a) IL4R^fl/fl^ CX3CR1Cre^+^ (Cre+) or IL4R^fl/fl^ CX3CR1Cre^-^(Cre^-^) mice were infected with 200 *T. muris* eggs and treated with αIFNγ or isotype mAb every 7 days during infection. Gut macrophages (gated as CD45^+^CD11b^+^CD11c^low/int^SiglecF^-^Ly6G^-^) were divided into P1 monocyte (CD64^-^Ly6C^hi^MHCII^-^), P2 transitioning monocyte (Ly6C^+^MHCII^+^) and P3 macrophage (CD64^+^Ly6C^-^MHCII^+^) compartments in the colon and caecum. (b) Representative plots illustrating the P1-3 gates in Cre^-^ and Cre^+^ mice following αIFNγ or isotype mAb at d18 post infection (c) Proportion of macrophages within the P1-P3 gates post infection with *T. muris.* Parasite-specific (d) IgG1 and (e) IgG2c production and (f) total IgE production at d18 post infection. Expression of the markers RELMα (g) and PD-L2 (h) within the total monocyte/macrophage (P1-3) population. Error bars show means ± SEM. Data are combined from two independent experiments (n=6-7). Statistical comparisons were performed with a two-way ANOVA with post-hoc Bonferroni’s multiple comparison test where appropriate: \**p*<0.05, \*\**p*<0.01, \*\*\**p*<0.001, \*\*\*\**p*< 0.0001 indicates significance compared to naïve control.

In conclusion, the pro- and anti-inflammatory properties of macrophages are linked to their activation state. Anti-inflammatory macrophage possess reparative functions, and thus significant interest exists in identifying how this activation state arises *in vivo.* Recent accounts report that anti-inflammatory macrophage activation can occur independently of IL4/IL13 during steady state in both the lung and the peritoneal cavity (5, 6), and that peritoneal macrophages and tumour associated macrophages can express RELMα and associated markers independently of IL4/IL13 during hypoxia or following lactate exposure. However, despite the physiological importance of the intestinal macrophage, a comprehensive analysis of the role of IL4Rα in driving the emergence of RELMα^+^Ym1^+^ macrophages has not been shown. This, in part, has been due to the limited tools to specifically target intestinal macrophages, with gene deletion with LysM cre poor in this compartment, especially during inflammatory conditions. Indeed, it is often inferred that these macrophages are IL4Rα dependent; the same is inferred to be true for the large intestine but has not been shown. Challenging prevailing paradigms, our data reveal a cell intrinsic IL4R alpha independence of the RELMα^+^Ym1^+^ large intestinal macrophage, even in highly polarised Th2 environments. We believe this to be an important discovery given that the large intestine is a physiologically relevant compartment for many inflammatory bowel disorders. Identifying the *in vivo* signals driving this anti-inflammatory macrophage phenotype is thus an urgent priority.

---

**Supplementary Figure 1.**
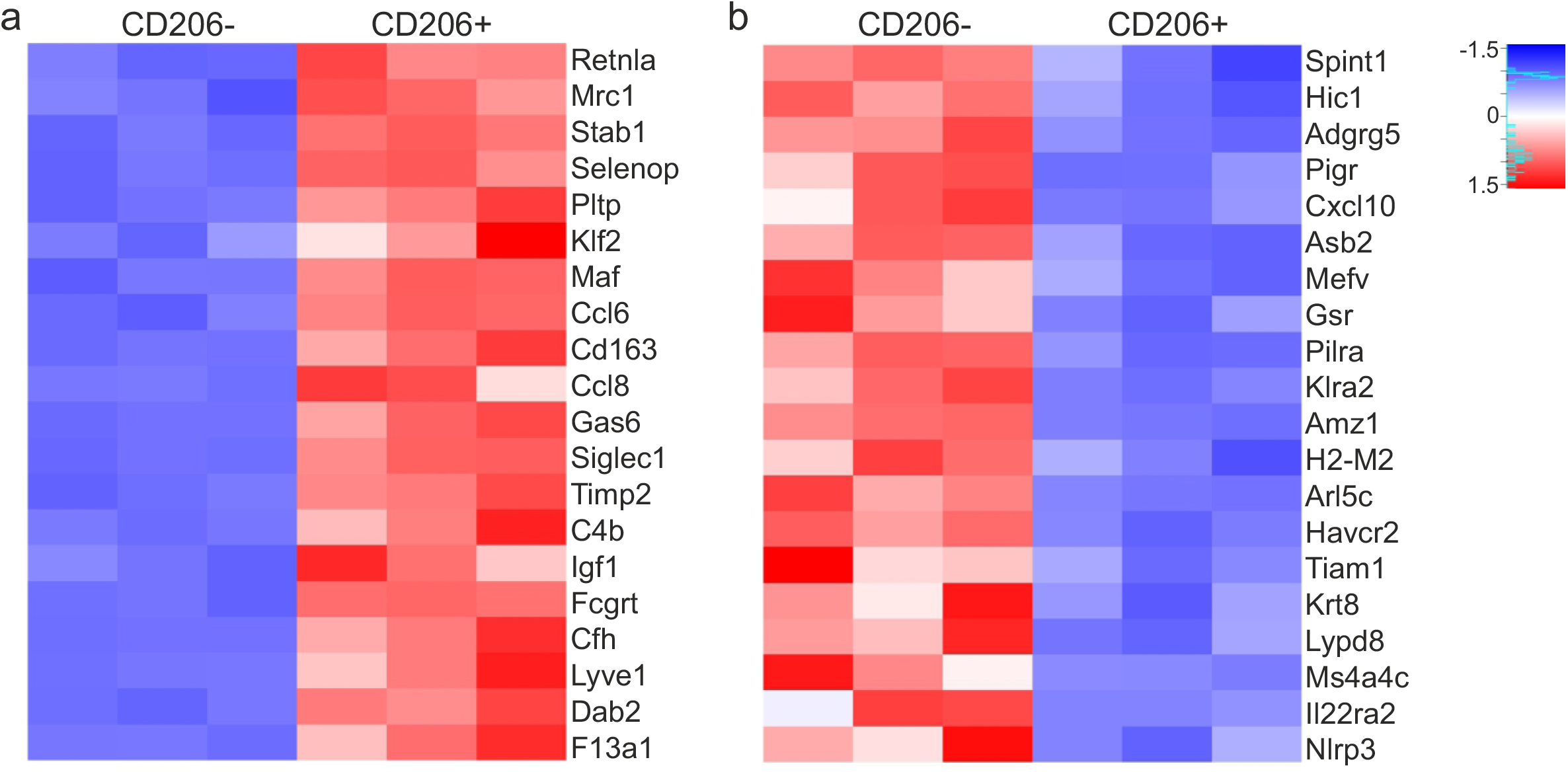
Confirmation of experimental models. Anti-inflammatory macrophages accumulate following *T. muris* infection. Gut macrophages (gated as CD45^+^CD11b^+^CD11c^low/int^SiglecF^-^Ly6G^-^) were FACS-sorted from infected C57 BL6/J mice at d35 post infection based on expression of CD206 and RNA-Seq performed. Analysis of (a) top 20 upregulated gene and (b) top 20 downregulated genes by copy number (blue indicates downregulation, red indicates upregulation).

**Supplementary Figure 2.**
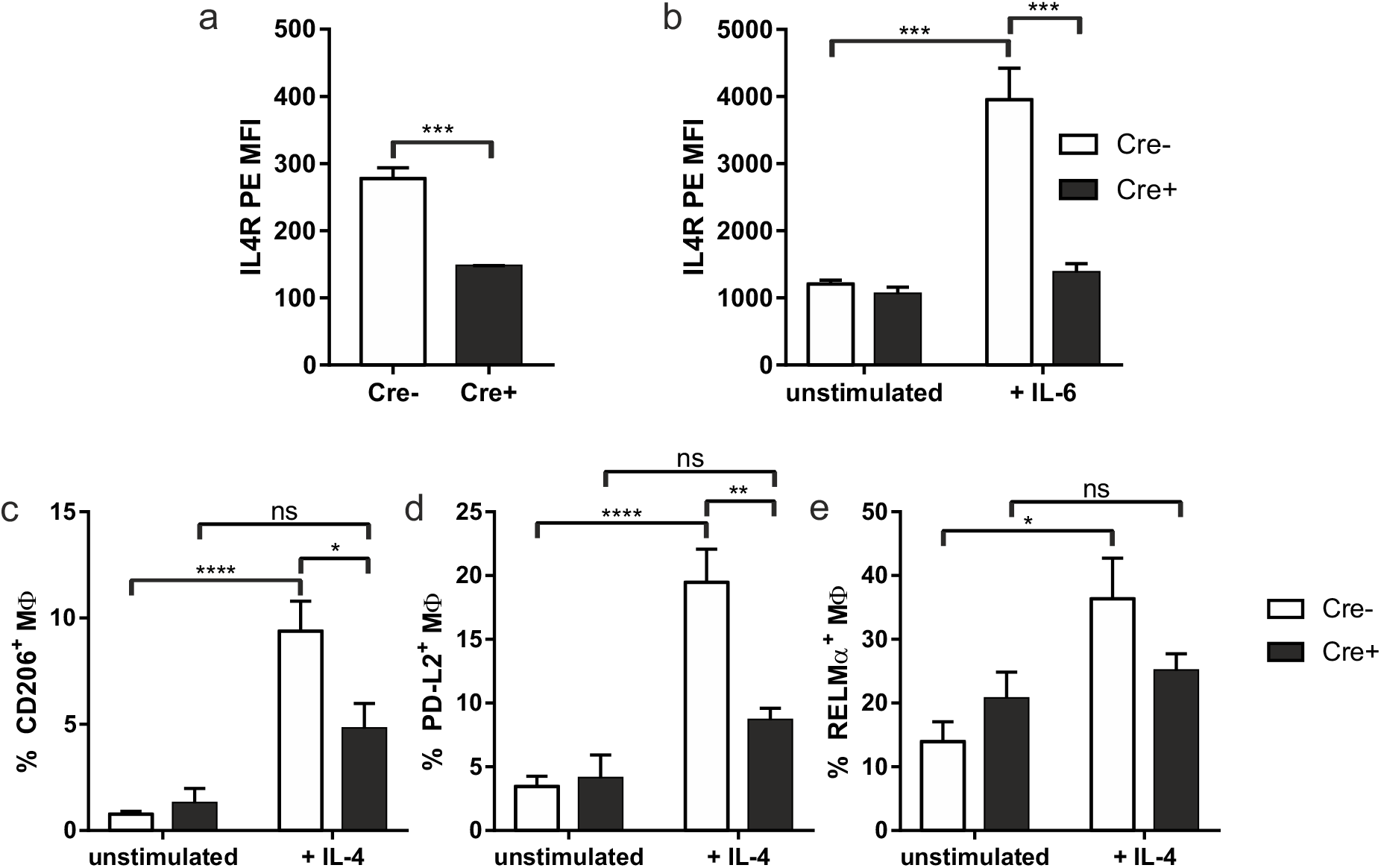
IL4R^fl/fl^ CX3CR1Cre^+^ macrophages are unresponsive to IL4 signalling. (a) IL4Rα expression was determined by flow cytometric analysis in gut macrophages at d35 post *T. muris* infection (n=3) or (b) Bone marrow derived macrophages (BMDMs) from Cre^+^ and Cre^-^ mice cultured for 18 hours in the presence or absence of IL6 for 18 hours. BMDMs were cultured in the presence of absence of IL4 for 24 hours and the expression of (c) CD206, (d) PD-L2 and (e) RELMα determined by flow cytometry. BMDM data is from 4-5 independent biological repeats. Error bars show means ± SEM. Statistical comparisons were performed with a two-way ANOVA with post-hoc Bonferroni’s multiple comparison test where applicable or by student’s *t*-test. Significance is shown compared to naïve control * *p*<0.05, \*\**p*<0.01, \*\*\**p*<0.001, \*\*\*\**p*< 0.0001 or as indicated.

**Supplementary Figure 3.**
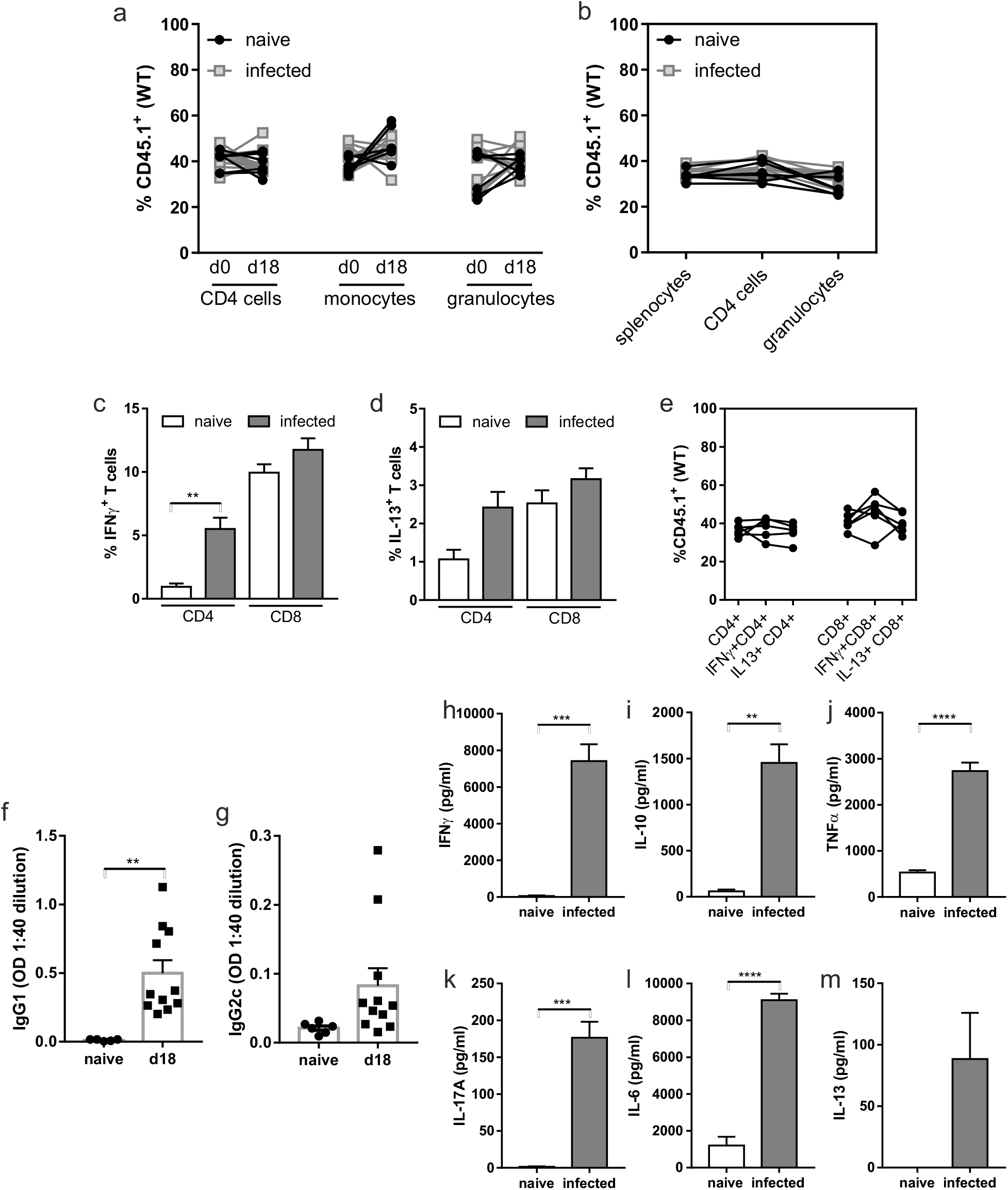
Mixed bone marrow chimeras show no survival advantage of IL4Rα expressing macrophages and mount a mixed Th1/Th2 response to *T. muris* infection. Mixed bone marrow chimeras were generated by lethally irradiating C57BL/6 IL4Rα^+/+^CD45.1^+^CD45.2^+^ mice and reconstitution with a 50:50 mix of IL4Rα^+/+^ CD45.1 and IL4Rα^-/-^ CD45.1^null^CD45.2^+/+^ congenic bone marrow cells; mice were left for 8 weeks to reconstitute. (a) Mice were tail bled prior to being left uninfected or infected with 200 *T. muris* eggs and blood chimerism analysed by assessing frequency of the CD45.1^+^ (WT) cells at d0 and d18 post infection, with lines joining individual mice. (b) Spleen populations analysed in naïve mice and at 18 post infection, with lines joining individual mice. CD4^+^and CD8^+^ T cells from the draining mesenteric lymph nodes were re-stimulated with eBioscience cell stimulation cocktail and intracellular cytokine expression of (c) IFNg and (d) IL13 analysed. (e) Frequency of the IL4Rα^+/+^CD45.1 cells within the different T cell populations was analysed at d18 post infection, with lines joining cells of individual mice. Parasite-specific (f) IgG1 and (g) IgG2c production in blood serum from naïve mice and at d18 post infection. MLN (h) IFNγ, (i) IL10, (j) TNF, (k) IL17A, (l) IL6 and (m) IL13 profiles in naïve mice and at d18 post infection cultured with *T. muris* E/S (50μg/ml) for 48h. Data are combined from two independent experiments (naïve n=6, d18 n=11). Error bars show means ± SEM. Statistical comparisons were performed as appropriate by repeated measure or standard two-way ANOVA with post-hoc Bonferroni’s multiple comparison test where applicable. Significance is shown compared to naïve control * *p*<0.05, \*\**p*<0.01, \*\*\**p*<0.001, \*\*\*\**p*< 0.0001 or as indicated.

## Detailed Methods

### Animals

All experimental procedures were approved by the University of Manchester Animal Welfare and Ethical Review Board and performed within the guidelines of the Animals (Scientific Procedures) Act, 1986. Male C57BL/6 mice were purchased from Envigo at age 6-8 weeks, CX3CR1 Cre mice were a gift from S. Jung. Severe combined immunodeficient (SCID) mice, IL4Rα^-/-^ mice, CD45.1^+^CD45.2^+^ mice, CD45.1^+/+^CD45.2^null^ and IL4Rαfl.CX3CR1Cre mice were bred in house at the University of Manchester and used at age 6-12 weeks. Mice were maintained at a temperature of 20-22°C in a 12h light, 12h dark lighting schedule, in sterile, individually ventilated cages with food and water *ad lib.* All IL4Rαfl.CX3CR1Cre mice used were littermate mice.

### *T. muris* passage

The parasite was maintained as previously described (45). Briefly, the parasite was passaged in susceptible SCID mice through infection of these mice with 150 infective *T. muris* eggs. At day 42 post infection the caecum and colon were removed, opened longitudinally, washed in pre-warmed RPMI-1640 media (Sigma-Aldrich, UK) supplemented with penicillin (Sigma-Aldrich; 500U/ml) and streptomycin (Sigma-Aldrich; 500μg/ml) (RPMI+5xP/S). Adult *T. muris* worms were gently removed using fine forceps under a dissecting microscope and carefully inspected to ensure they retained no epithelial cells. Worms were then cultured in 6 well tissue culture plates containing RPMI + 5x P/S. Plates were incubated in a moist humidity box for 4 hours at 37°C to collect 4h E/S then each well split into two wells containing fresh medium and incubated again in a humidity box at 37°C overnight. Media from both the 4hr culture and the overnight culture were processed in the same way to retrieve unembryonated eggs and E/S products.

### Preparation of Eggs

Media from worm cultures was centrifuged at 720g for 15 mins at room temperature. Supernatant was removed and kept for preparation of E/S. Pelleted eggs were resuspended in 40 deionised water, filtered through a 100μm nylon sieve and transferred to a cell culture flask. Flasks were examined under a dissecting microscope (Leica S8 APO) and egg density adjusted as necessary. To allow embryonation to occur, flasks were wrapped in foil to ensure eggs were kept in darkness, stored horizontally and eggs were monitored weekly. If any fungal growth was observed eggs were re-filtered or washed by centrifugation and transferred to fresh flasks. After approximately 8 weeks eggs were fully embryonated and transferred for storage, horizontally at 4°C. In order to determine the number of eggs required to establish around 150 worms *in vivo,* all egg batches were tested in SCID mice prior to experimental use to determine the infectivity of each batch of eggs. A minimum of 3 SCID mice were infected with approximately 200 eggs and the number of larvae present in the colon and caecum at day 14 post infection determined. The infectivity was then calculated as the % larvae counted from a given number of eggs. For high-dose *T. muris* infections approximately 3ml of egg suspension was transferred to a universal tube, 15ml of deionisied water added prior to centrifugation for 15mins at 720g (low brake), room temperature. Pelleted eggs were resuspended in MQ water and embryonated eggs counted adjusted, in accordance with calculated infectivity data, to ensure there were 150 infective eggs per 200ul. Mice were then infected with 150 infective T. muris eggs in 200ul by oral gavage. During both egg counting and infection eggs were prevented from settling by stirring on a magnetic stirrer.

### Preparation of excretory/secretory products

4hr and overnight supernatants containing E/S products from passage worm cultures were filter sterilised through a 0.2μm syringe filter (Merck, Germany). E/S was then concentrated using Amicon Ultra-15 centrifugal filter units (Merck) by centrifugation at 3,000g at 4°C. Centrifugation was carried out until approximately 20% of the original volume of supernatant remained. E/S was then dialysed against PBS using Slide-A-Lyzer Dialysis cassettes, 3.500 MWCO (Thermo Scientific, UK) at 4°C. The E/S was then filtered sterilised a final time through a 0.2μm syringe filter, the protein concentration measured and aliquoted prior to storage at −80°C

### Quantification of *T. muris* worms

At necropsy the caecum and proximal colon were collected, blinded and stored at −20°C. For worm counts, frozen caecum and colon were defrosted in a petri dish containing water and cut open longitudinally. The gut contents were removed through swilling in the water and the gut tissue transferred to a fresh petri dish containing water. The gut mucosa was scraped off using curved forceps to remove epithelia and worms from the gut tissue. Both the gut contents and removed gut mucosa were then examined and *T. muris* worms counted using a dissecting microscope (Leica S8 APO).

### *In vivo* antibody treatment

Monoclonal antibodies against IFNγ(XMG-1.2) and control isotype (GL113) were a gift from L. Boon. Mice were treated intraperitoneally with 1 mg of antibody on day 0, day 7 and day 14 post infection.

### Bone Marrow Chimeras

Competitive mixed BM chimeric mice were created by lethally irradiating C57BL/6 CD45.1^+^CD45.2^+^ mice with 11.5 Gy γ radiation administered in two doses ~3 h apart, followed by i.v. injection of 9 × 10^6^ BM cells depleted of mature T cells using CD90 microbeads (Miltenyi Biotec) and comprised of a 1:1 mix of cells from C57BL/6 CD45.2^+/+^IL4Rα^-/-^ mice and C57BL/6 CD45.1^+/+^CD45.2^null^ mice. Chimeric animals were left 8 weeks before further experimental manipulation.

### Tissue preparation and cell isolation

#### Caecum and colon lamina propria

Cells were isolated as previously described with some modifications (46). In brief, after dissection of the caecum and colon, the colonic patch was removed from the tip of the caecum and all adherent adipose tissue removed from the tissue. The caecum and colon were cut longitudinally and washed thoroughly with pre-chilled PBS on ice. To remove intestinal epithelial cells and leukocytes, the tissue was incubated in prewarmed media (RPMI 1640) supplemented with 3% FCS (Sigma-Aldrich), 20 mM Hepes (Sigma-Aldrich), 5 mM EDTA (VWR, UK), and 1 mM freshly thawed dithiothreitol (Sigma-Aldrich) for 20 min at 37°C with agitation. After incubation, gut segments were shaken three times in fresh prewarmed RPMI 1640 (serum-free) with 2 mM EDTA and 20 mM Hepes. The remaining tissue was minced with fine scissors and digested at 37°C for 30 min with continuous stirring at 450 rpm in serum-free RPMI 1640 containing 20 mM Hepes, 0.1 mg/ml liberase TL (Sigma-Aldrich), and 0.5 mg/ml DNase I (Sigma-Aldrich). Digested tissue was passed sequentially through a 70-μm filter and 40-μm cell strainer, and after pelleting, it was resuspended in media supplemented with 10% FCS until staining.

#### Blood

Blood was collected into heparin-coated 0.2mm capillary tubes (VWR) and stored in EDTA (VWR). Red blood cells were lysed in RBC lysis buffer (Sigma-Aldrich) for 5 min on ice twice. Cells were then washed and resuspended in PBS containing 10% FCS until staining.

#### Culture of bone marrow macrophage cells

Bone marrow-derived macrophages (BMDM) were isolated from mice by flushing femurs with Dulbecco’s modified Eagle’s medium (DMEM; Invitrogen) containing 10 % FCS, 1 % L-glutamine and 100U/ml penicillin/100μg/ml streptomycin (all from Sigma-Aldrich, UK) (complete DMEM). Cells were washed and cultured at 1 × 10^6^/ml in complete DMEM containing 30 ng/ml M-CSF (Peprotech, UK) for 7 days, with the media being replaced after 4-5 days. Macrophage purity was assessed by flow cytometry and in all cultures used the purity was found to be over 90%. The BMDMs (0.5 × 10^6^/ml) were stimulated for 24 hours with IL4 (20ng/ml; Peprotech) or IFNγ (10ng/ml; eBioscience, UK) and LPS (10ng/ml; Sigma-Aldrich) or for 18 hours with IL6 (50ng/ml; Peprotech). All supernatants were stored at −20 °C until analysed, cells were removed using Accutase (Thermo Scientific) prior to flow analysis.

### Flow cytometry

Single-cell suspensions of caecum/colon, blood, bone marrow derived macrophages, MLN or spleen were washed thoroughly with PBS and stained with the Live/Dead Fixable blue dead cell stain kit (Thermo Scientific) in the dark for 15 min at 4°C to exclude dead cells. Subsequently, cells were stained in the dark for 10 min at 4°C with anti-CD16/CD32 (eBioscience) in PBS containing 0.5% BSA (Sigma-Aldrich) and a further 20min with the relevant fluorochrome-conjugated antibodies in Brilliant stain buffer (BD biosciences, UK). Cells were washed and in some cases, cells were immediately acquired live, or alternatively, cells were fixed in fixation buffer (True-Nuclear Transcription factor buffer set, Biolegend, UK) for 20 min at 4°C and resuspended in PBS containing 0.5% BSA. For intracellular staining cells were permeabilised (True-Nuclear Transcription factor buffer set, Biolegend) and stained in permeabilisation buffer with the relevant fluorochrome-, biotin-conjugated or purified antibodies for 20 min at 4°C. Cells were washed in permeabilisation buffer and resuspended when relevant with fluorochrome-conjugated secondary antibodies and then washed in PBS prior to resuspending in PBS containing 0.5% BSA for acquisition. Cell acquisition was performed on an LSR Fortessa or LSR II running FACSDIVA 8 software (BD biosciences). For each intestinal sample, typically 10,000-20,000 macrophages were collected. Data were analysed using FlowJo software version 10 (TreeStar, UK).

### Gut monocyte and macrophage isolation

Single-cell suspensions were prepared as described above with the following modifications: incubation with dithiothreitol and EDTA was reduced to 10 min, and the liberase digestion step was decreased to 20 min but with an increased concentration of liberase TL (0.75 mg/ml). Cells were sorted on an Influx (BD biosciences) and isolated cells were suspended in PBS supplemented with 2% FCS (Sigma-Aldrich) and 2 mM EDTA (VWR). Sorted cells were collected in PBS with 10% FCS and stored on ice prior to washing and resuspending in Trizol (Thermo Fisher) before storage at −80°C for subsequent RNA extraction.

### RNA extraction

RNA was extracted from sorted cells using an RNeasy mini kit (QIAGEN, UK) following the manufacturer’s instructions.

### RNA sequencing and analysis

Total RNA was submitted to the University of Manchester Genomic Technologies Core Facility. Quality and integrity of the RNA samples were assessed using a 2200 TapeStation (Agilent Technologies) and then libraries generated using the TruSeq^®^ Stranded mRNA assay (Illumina, Inc.) according to the manufacturer’s protocol. Briefly, total RNA was used as input material from which polyadenylated mRNA was purified using poly-T, oligo-attached, magnetic beads. The mRNA was then fragmented using divalent cations under elevated temperature and then reverse transcribed into first strand cDNA using random primers. Second strand cDNA was then synthesised using DNA Polymerase I and RNase H. Following a single ‘A’ base addition, adapters were ligated to the cDNA fragments, and the products then purified and enriched by PCR to create the final cDNA library. Adapter indices were used to multiplex libraries, which were pooled prior to cluster generation using a cBot instrument. The loaded flow-cell was then paired-end sequenced (76 + 76 cycles, plus indices) on an Illumina HiSeq4000 instrument. Finally, the output data was demultiplexed (allowing one mismatch) and BCL-to-Fastq conversion performed using Illumina’s bcl2fastq software, version 2.17.1.14

Unmapped paired-end sequences from an Illumina HiSeq4000 sequencer were tested by FastQC (http://www.bioinformatics.babraham.ac.uk/projects/fastqc/). Sequence adapters were removed and reads were quality trimmed using Trimmomatic_0.36 (PMID: 24695404). The reads were mapped against the reference mouse genome (mm10/GRCm38) and counts per gene were calculated using annotation from GENCODE M14 (http://www.gencodegenes.org/) using STAR_2.5.3 (PMID: 23104886). Normalisation, Principal Components Analysis, and differential expression was calculated with DESeq2_1.16.1 (PMID:25516281).

### Mesenteric lymph node (MLN) cell re-stimulation and cytokine bead array

MLN cells were brought to cell suspension and 5×10^6^ cells/ml were cultured for 48h at 37°C 5% CO2 in RPMI 1640 with 4 hr E/S antigen (50μg/ml). Supernatants were harvested and stored at −20°C until they were assayed for cytokines. Cytokines were analysed using the Cytometric Bead Array (CBA) Mouse/Rat soluble protein flex set system (BD Bioscience), which was used according to the manufacturer’s instructions. Cell acquisition was performed on a FACS Verse (BD Biosciences) or MACSQuant (Miltenyi Biotech). For analysis, FCAP Array v3.0.1 software (BD Cytometric Bead Array) was used. For intracellular cytokine analysis MLN cells were brought to cell suspension and 5×10^6^cells/ml were cultured for 16h at 37°C 5% CO_2_ with eBioscience Cell Stimulation cocktail (Thermo Fisher Scientific) prior to flow cytometric analysis.

### Histology

Proximal colon tissue was fixed for 24 hours in 10% neutral buffered formalin (Fisher) containing 0.9% sodium chloride (Sigma-Aldrich), 2% glacial acetic acid (Sigma-Aldrich) and 0.05%alkyltrimethylammonium bromide (Sigma-Aldrich) prior to storage in 70% Ethanol (Fisher) until processing. Fixed tissues were dehydrated through a graded series of ethanol, cleared in xylol and infiltrated with paraffin in a dehydration automat (Leica ASP300 S) using a standard protocol. Specimens were embedded in paraffin (Histocentre2, Shandon), sectioned on a microtome (5μm sections) and allowed to dry for a minimum of 4 hours at 40°C. Prior to staining slides were deparaffinised with citroclear (TCS biosciences) and rehydrated through alcohol (100% to 70%) to PBS or water.

Mucins in goblet cells were stained with 1% alcian blue (Sigma-Aldrich) in 3% acetic acid (Sigma-Aldrich, pH 2.5) for 5 mins. Sections were washed and treated with 1% periodic acid, 5mins (Sigma-Aldrich). Following washing sections were treated with Schiff’s reagent (Vicker’s Laboratories) for 15mins and counterstained in Mayer’s haematoxylin (Sigma-Aldrich). Slides were then dehydrated and mounted in depex mounting medium (BDH Laboratory Supplies). For enumeration of goblet cell staining, the average number of cells from 20 crypts was taken from three different sections per mouse. Images were acquired on a 3D-Histech Pannoramic-250 microscope slide-scanner using a *20x/0.30 Plan Achromat* objective (Zeiss).

### Immunohistochemistry

Proximal colon tissue was embedded in OCT and snap-frozen on dry ice and stored at −80C until processing. Cryostat frozen sections (6 μm) were fixed in acetone, blocked with blocking solution (Perkin-Elmer) plus 7% goat serum (Vector) prior to incubation with 2 μg/mL rabbit anti-mouse RELMa (Peprotech) followed by anti-rabbit Alexa488 (2.5 μg/ml, Invitrogen). Sections were re-blocked with blocking solution (Perkin-Elmer) plus 7% rat serum (Vector) prior to incubation with rat anti-mouse CD68-AF647 (1 μg/ml, BioLegend). Sections were counterstained and mounted with VectaShield Hard set mounting media with DAPI (Vector). Images were collected on a Zeiss Axioimager D2 upright microscope using a 20x / 0.5 EC Plan Apochromat objective and captured using a Coolsnap HQ2 camera (Photometrics) through Micromanager software v1.4.23. Specific band pass filter sets for DAPI, FITC and Cy5 were used to prevent bleed through from one channel to the next. Images were then processed and analysed using *Fiji ImageJ (http://imagej.net/Fiji/Downloads)*.

### IgG ELISA

Serum was assayed for parasite specific IgG1 and IgG2c antibody production. 96 well plates were coated with 5μg/ml *T. muris* overnight E/S antigen overnight, plates were washed, and non-specific binding blocked with 3% BSA (Sigma-Aldrich) in PBS. Following washing, plates were incubated with serum (2 fold dilutions, 1:20-1:2560) and parasite specific antibody was measured using biotinylated IgG1 (BD Biosciences) or IgG2c (BD Biosciences) antibodies which were detected with SA-POD (Roche). Finally, plates were washed and developed with TMB substrate kit (BD Biosciences, Oxford, UK) according to the manufacturer’s instructions. The reaction was stopped using 0.18 M H2SO4, when sufficient colour had developed. The plates were read by a Versa max microplate reader (Molecular Devices) through SoftMax Pro 6.4.2. software at 450 nm, with reference of 570 nm subtracted.

### IgE ELISA

Serum was assayed for total IgE antibody production. 96 well plates were coated with purified antimouse IgE (2ug/ml, Biolegend, Clone: RME-1) in 0.05M carbonate/ bicarbonate buffer and incubated overnight at 4°C. Following coating, plates were washed in PBS-Tw and non-specific binding blocked with 3% BSA (Sigma-Aldrich) in PBS for 1 hour at room temperature. Plates were washed and diluted serum (1:10) added to the plate and incubated for 2hrs at 37°C. After washing HRP conjugated goat anti-mouse IgE (1ug/ml; Bio-rad) was added to the plates for 1 hour. Finally, plates were washed and developed with TMB substrate kit (BD Biosciences, Oxford, UK) according to the manufacturer’s instructions. The reaction was stopped using 0.18M H2SO4, when sufficient colour had developed. The plates were read by a Versa max microplate reader (Molecular Devices) through SoftMax Pro 6.4.2. software at 450 nm, with reference of 570 nm subtracted.

### Statistical analysis

Comparisons between groups were undertaken using Prism (8.0; GraphPad Software). Two experimental groups were compared using a Student’s *t* test, where more than two groups were compared, a one-way ANOVA or two-way ANOVA was used as appropriate. Significance was set at *p*≤0.05.

### Flow cytometry antibodies used

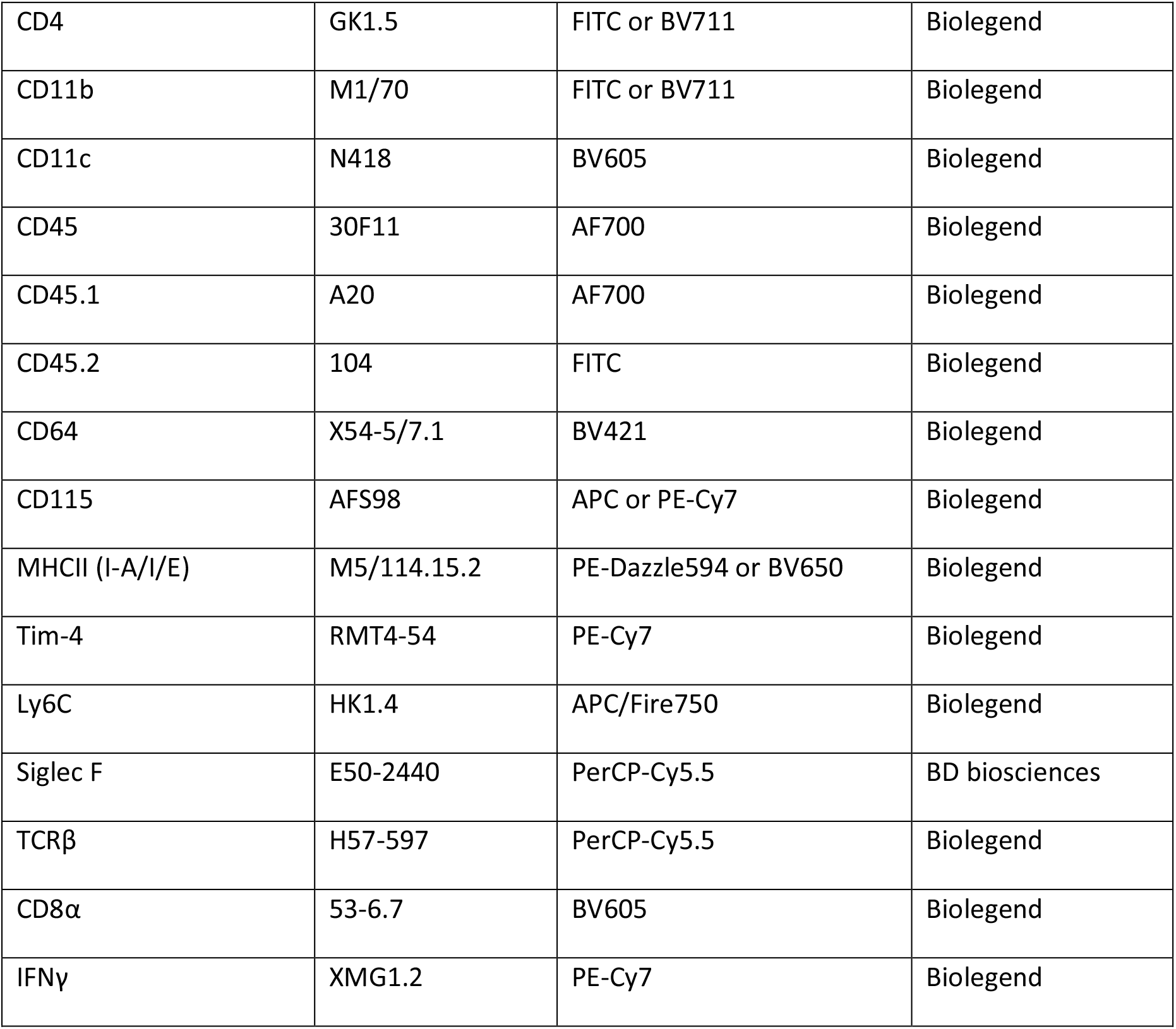
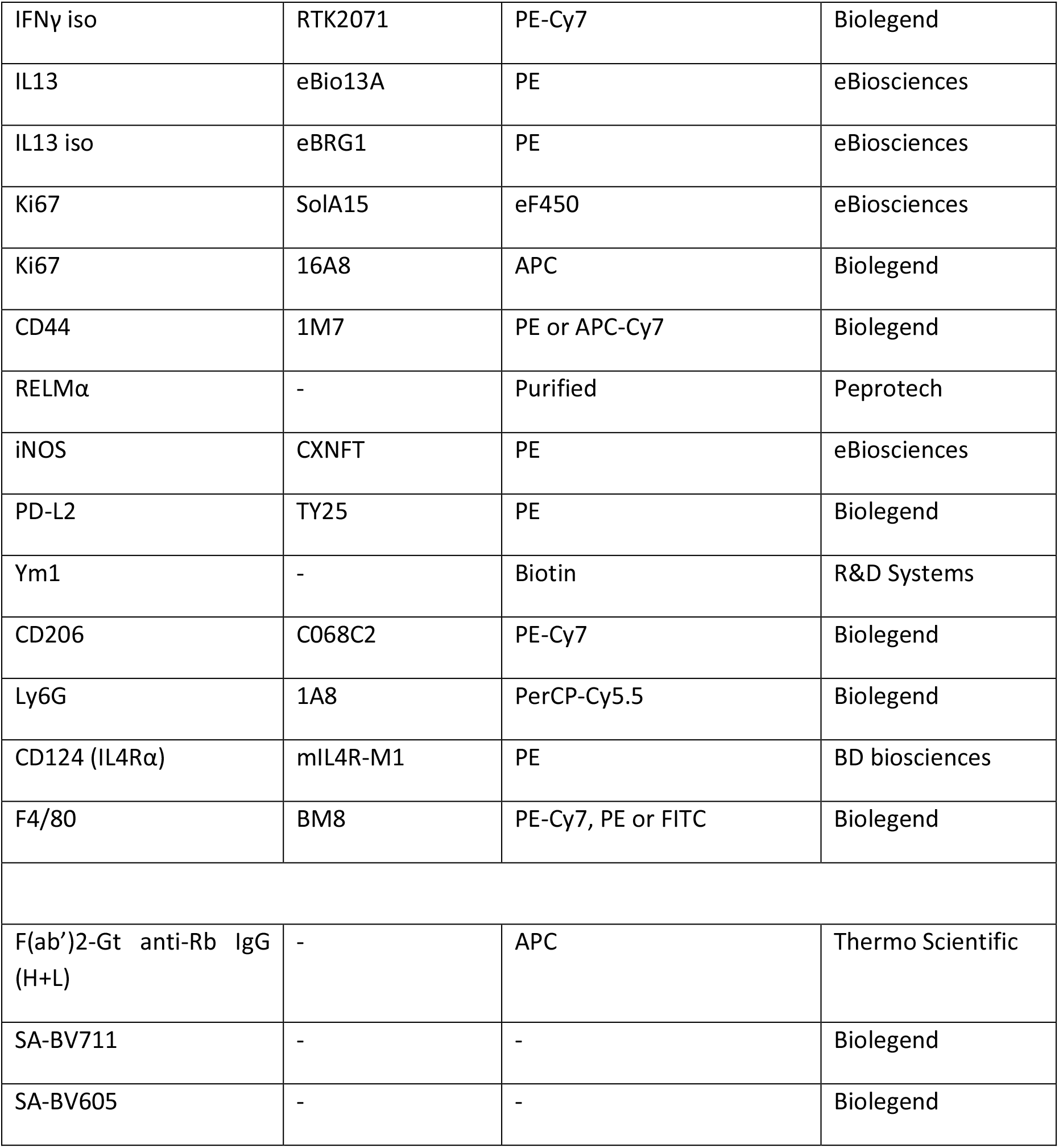

## Author contributions

R Forman conceptualized the studies, designed experiments, performed research, analyzed and interpreted the data, and wrote the manuscript; L. Logunova provided expertise, developed methodologies and performed experiments; H. Smith, K. Wemyss, I. Mair provided expertise, performed experiments, and edited the manuscript; J.E. Allen, W. Muller and J.L. Pennock provided expertise in interpreting the data, contextualizing the studies, and editing the manuscript; K. J. Else conceptualized the studies, supported design of experiments, analysis, and interpretation of data, and wrote the manuscript.

## Acknowledgements

Thanks goes to the following core facilities at The University of Manchester: the bioimaging, histology, flow cytometry, genomic technologies, bioinformatics, and biological services facilities. The Bioimaging Facility microscopes used in this study were purchased with grants from BBSRC, Wellcome and the University of Manchester Strategic Fund. The Histology Facility equipment used in this study was purchased with grants from the University of Manchester Strategic Fund. Irradiation in these experiments was performed with assistance from Epistem Ltd and Dr Joanne Konkel. Special thanks goes to Roger Meadows, Steve Marsden, Gareth Howell, Matthew Brown, Leo Zeef and Andy Hayes for their technical assistance. We thank Dr Steve Jenkins, Dr Tara Sutherland, Dr John Grainger and Dr Dominick Ruckerl for their discussions of the data and experimental design.

This work was supported by a MRC grant (MR/N022661/1) awarded to K.J. Else.

The authors declare no competing financial interests.

